# Protectin D1/GPR37 signaling enhances macrophage-dependent efferocytosis to attenuate experimental abdominal aortic aneurysm formation

**DOI:** 10.1101/2025.10.20.683538

**Authors:** Aravinthan Adithan, Michael Fassler, Guanyi Lu, Jeff Arni C. Valisno, Gang Su, Shiven Sharma, Walker Ueland, Ashish K. Sharma, Gilbert R. Upchurch

## Abstract

Abdominal aortic aneurysms (AAAs) are chronic inflammatory vascular disorders characterized by progressive aortic dilation and destruction of the vascular wall, often culminating in rupture. Current management is limited to surgical repair, with no approved targeted pharmacologic therapies. In this study, we investigated the immunomodulatory role of Protectin D1 (PD1), a specialized pro-resolving lipid mediator, through G-protein–coupled receptor 37 (GPR37) signaling on macrophages in mitigating AAA progression and preventing aortic rupture. Single cell-RNA sequencing analysis of human tissue demonstrated significant differences in PD1/GPR37 axis-related genes in macrophages in AAAs compared to control aortic tissue. Using an established murine AAA model, PD1 administration significantly attenuated aortic diameter, pro-inflammatory cytokine and matrix metalloproteinase (MMP2) expression, as well as maintained aortic morphology in a GPR37-dependent manner. Importantly, PD1 treatment prevented preformed AAA progression to aortic rupture in another preclinical elastase+BAPN model of aortic rupture, by attenuating aortic diameter, tissue inflammation as well as decreasing macrophage infiltration, preserving elastin integrity, and restoring smooth muscle α-actin expression in the aortic wall. Mechanistically, PD1 enhanced macrophage efferocytosis of apoptotic vascular smooth muscle cells in the murine aortic tissue as well as in isolated macrophages via GPR37-dependent manner and attenuated the inflammatory paracrine secretion of macrophage-specific paracrine release of TNF-α and IL-β. These findings suggest that PD1/GPR37 signaling on macrophages promotes inflammation-resolution by enhancing efferocytosis of apoptotic SMCs conferring protection against aortic inflammation and remodeling to mitigate AAA formation and rupture.

**Significance Statement:** This study elucidates the protective role of specialized proresolving lipid mediator, Protectin D1, by activating macrophages via GPR37 receptors, to enhance the clearance of apoptotic smooth muscle cells to mitigate aortic inflammation and vascular remodeling during abdominal aortic aneurysm formation. We observed that several key inflammation related genes were associated with PD1/GPR37-dependent signaling in macrophages of human AAAs. Detailed analysis in experimental models delineated the signaling pathway where upregulating macrophage-dependent efferocytosis via immunomodulation by Protectin D1 attenuated aortic inflammation and remodeling, indicating a potential mechanism for therapeutic intervention in the pathobiology of AAAs to prevent aortic rupture.

## INTRODUCTION

Abdominal aortic aneurysm (AAA) formation is a progressive and potentially fatal vascular disorder characterized by the localized dilation of the abdominal aorta, which increases the risk of rupture(1). The global burden of AAA continues to rise, particularly in aging populations, with high morbidity and mortality rates due to the lack of effective pharmacological interventions. Current treatment strategies rely primarily on surgical repair, such as open surgery or endovascular aneurysm repair, but these approaches are invasive and associated with complications, particularly in high-risk patients(2). Thus, there is an urgent need for novel therapeutic interventions targeting the underlying mechanisms of AAA progression.

The pathogenesis of AAA is complex and multifactorial, driven by chronic inflammation, extracellular matrix degradation, thrombus formation, and the accumulation of apoptotic cell debris within the aortic wall(3). These pathological processes result in immune dysregulation, promoting macrophage infiltration, increased oxidative stress, and the release of proteolytic enzymes such as matrix metalloproteinases (MMPs), which degrade the structural integrity of the aortic wall(4). Additionally, inefficient clearance of apoptotic cells further perpetuates a pro-inflammatory microenvironment, accelerating tissue destruction and aneurysm expansion(5).

Efferocytosis, the process by which phagocytic cells such as macrophages recognize and engulf apoptotic cells, is a fundamental mechanism in maintaining tissue homeostasis and resolving inflammation(6). In the context of AAA, defective macrophage efferocytosis results in the accumulation of necrotic and apoptotic debris, sustaining chronic inflammation and exacerbating vascular remodeling(7). Emerging evidence suggests that specialized pro-resolving mediators (SPMs), such as Protectin D1 (PD1), play a pivotal role in enhancing macrophage-dependent efferocytosis, facilitating the resolution of inflammation, and preventing further vascular damage(8). PD1, a bioactive lipid mediator derived from docosahexaenoic acid (DHA), has been shown to exert potent anti-inflammatory and pro-resolving effects by modulating immune cell function(9). Recent studies highlight that PD1 exerts its protective effects through G-protein-coupled receptor 37 (GPR37) signaling, a receptor implicated in macrophage activation and apoptotic cell clearance but the role of PD1/GPR37 remains to be elucidated(10). The binding of PD1 to GPR37 initiates intracellular signaling cascades that enhance macrophage efferocytosis, leading to the effective clearance of apoptotic debris and suppression of pro-inflammatory cytokine production(11). In addition to its role in efferocytosis, PD1/GPR37 signaling has been shown to modulate key inflammatory pathways involved in vascular diseases, including NF-κB and MAPK signaling, which are central regulators of inflammatory cytokine expression and immune cell recruitment(12). Since AAA is driven by sustained inflammation and matrix degradation, our hypothesis focused on delineating the mechanistic role of PD1/GPR37 activation on macrophages to mitigate downstream aortic inflammation and vascular remodeling during AAA formation and aortic rupture.

Our data suggests that PD1/GPR37-dependent genes are differentially regulated during human AAA formation. Furthermore, treatment with PD1 significantly attenuates aortic diameter, proinflammatory cytokine expression, enhances apoptotic cell clearance and restores vascular integrity to decrease AAA formation and prevent aortic rupture. In summary, the PD1/GPR37 signaling pathway offers a novel and promising approach for attenuating AAA progression by enhancing macrophage-dependent efferocytosis and resolving inflammation. By reducing apoptotic cell accumulation and modulating immune responses, PD1 may provide a targeted pharmacological intervention for AAA, addressing the critical gap in current treatment options.

## MATERIALS AND METHODS

### Animals

Male wild-type C57BL/6 and GPR37^-/-^ mice, aged 8 to 12 weeks, were obtained from Jackson Laboratory (Bar Harbor, ME). The animals were housed under controlled conditions—70°F temperature, 50% humidity, and a 12-hour light–dark cycle—according to institutional animal care guidelines. Mice had free access to drinking water and a standard chow diet. All experimental procedures were approved by and conducted in accordance with the Institutional Animal Care and Use Committee of the university of Florida (protocol #201910902).

### Human Single Cell RNA Sequencing

Single cell RNA-seq data set of human AAAs and controls was obtained from Gene Expression Omnibus (GSE166676) and re-analyzed via Seurat(13). Cell annotations were assigned by extracting cluster specific markers via FindMarkers and cross-referencing the marker set with published human and mouse cell atlases via PanglaoDB(14). Cell annotations were verified via expression of putative cell markers for macrophage clusters as described in previously published analyses of scRNA datasets of the aorta(15, 16). Differential gene expression in macrophage clusters between human AAA and controls were performed via the FindMarkers function in Seurat and statistical significance was assessed using the Wilcoxon Rank Sum Test. Differentially expressed genes (DEGs) were identified as genes with adjusted p-value < 0.05 and |fold change| >0.1 A list of PD1 and GPR37-receptor related genes were generated using GeneCards (query=Protectin D1 or GPR37) and genes with relevance score >1 were cross-references with list of DEGs to identify differentially expressed PD1 and GPR37 receptor related genes (DE-PGRGs)..

### Murine AAA models

Mice were anesthetized using isoflurane, and the infrarenal abdominal aorta was exposed through a midline laparotomy, as previously described(17). The aorta was carefully dissected circumferentially from surrounding tissues and topically treated for 5 minutes with 5 μL of elastase (0.4 U/mL, type I porcine pancreatic elastase; Sigma-Aldrich, St. Louis, MO) or heat-inactivated elastase (HIE), which served as a control. In the AAA and aortic rupture model, elastase application was combined with supplementation of 0.2% β-aminopropionitrile (BAPN) in drinking water, continued through day 28 post-surgery. Treated aortic sections were harvested post operatively on day 14 (topical elastase model) and day 28 (AAA and aortic rupture model), and preserved in paraformaldehyde for immune histochemistry or snap-frozen in liquid nitrogen for molecular analysis. Aortic diameter was measured using video micrometry (AmScope, Irvine, CA). The degree of aortic dilation was calculated using the formula: [(maximal aortic diameter − baseline diameter) / baseline diameter] × 100. A dilation of ≥100% was considered indicative of AAA formation.

### Histology

Harvested aortic tissues were fixed overnight in 10% zinc-buffered formalin, followed by dehydration through a graded ethanol series and embedding in paraffin. For immunohistochemical analysis, tissue sections were stained using the following primary antibodies: anti-mouse Mac-2 for macrophages (1:5000; Cedarlane Laboratories, Burlington, ON, Canada; catalog no. CL8942AP) and anti-mouse α-smooth muscle actin (α-SMA, 1:1000; Sigma, St. Louis, MO; catalog no. A5691). Additional staining included Verhoeff–Van Gieson (VVG) stain for elastin (Polysciences, Inc., Warrington, PA; catalog no. 25089-1). Elastin degradation was quantified by counting the number of elastin fiber breaks per mm² using Fiji (ImageJ, NIH), and results were

averaged and presented graphically. Histological evaluation was performed on three separate aortic sections per animal, and quantification was conducted independently by blinded observers. Representative images from multiple animals per group were selected for consistency and accuracy. Imaging was performed at 20× magnification using a Nikon microscope equipped with a digital camera and NIS-Elements BR software. Quantification of staining intensity was performed using QuPath version 0.5 (QuPath; University of Edinburgh, Edinburgh, UK) by measuring the percentage of positively stained area relative to the total aortic cross-sectional area.

### *In vitro* efferocytosis assay

Efferocytosis was evaluated using thioglycollate-elicited mouse peritoneal macrophages and apoptotic MOVAS (mouse aortic smooth muscle cells; ATCC, Manassas, VA) cells as targets. Male C57BL/6 or GPR37^-/-^ mice were injected intraperitoneally with 3% Brewer thioglycollate broth (Sigma-Aldrich, St. Louis, MO). After 72 hours, peritoneal exudate cells were collected via lavage with ice-cold sterile PBS. The harvested cells were cultured in RPMI-1640 medium supplemented with 10% fetal bovine serum (FBS) and 1% penicillin-streptomycin and incubated at 37°C in a 5% CO₂ atmosphere for 2 hours. The adherent macrophages were pretreated with or without PD1 (50 nM) for 2 hrs prior to the assay. MOVAS cells were labeled with PKH67 green-fluorescent dye (Sigma-Aldrich) following the manufacturer’s protocol. Apoptosis was induced by treating the labeled cells with 1 μM staurosporine (Fisher Scientific) for 3 hours at 37°C. After washing, apoptotic MOVAS cells were co-cultured with peritoneal macrophages at a 1:1 ratio (1 × 10⁶ cells each) in 6-well plates and incubated for 45 minutes at 37°C. Following incubation, unengulfed apoptotic cells were removed by PBS washing. Cells were then stained for Live/Dead staining (Thermo Fisher Scientific), followed by PE-Cy7-conjugated anti-mouse F4/80 antibody (BioLegend, San Diego, CA) to identify macrophages. Flow cytometry was used to quantify efferocytosis, defined as the percentage of F4/80⁺ macrophages also positive for PKH67 fluorescence, indicating internalization of apoptotic MOVAS cells(18).

### *In vivo* efferocytosis Assay

Mice in the study group received 2 intraperitoneal injection of PD1 (300 ng/mouse) on day 1 and 2. On day 3, murine abdominal aortas were excised under sterile conditions, cleaned of periadventitial tissue, and placed in cold DMEM in the elastase-treatment model. Tissues were minced into ∼1 mm³ fragments and digested in type I collagenase (2.5 mg/mL, Worthington Biochemical) and porcine pancreatic elastase (0.5 mg/mL, Sigma-Aldrich) prepared in DMEM plain medium at 37 °C for 40 min with gentle agitation. Digests were passed through a 70-µm strainer and collected into ice-cold staining buffer (PBS with 2% FBS and 2 mM EDTA). Cell suspensions were centrifuged at 400 × g for 5 min at 4 °C and washed prior to Live/Dead blue fixable staining as per the manufacturer’s instruction staining. Cells were incubated with anti-mouse CD16/32 (Fc Block, BioLegend, 10 µg/mL) for 10 min at 4 °C to minimize nonspecific binding and then stained for 30 min at 4 °C in the dark with anti-mouse CD45-BV785 (BioLegend) and anti-mouse F4/80-PE-Cy7 (eBioscience). Cells were fixed and permeabilized using BD Perm/Wash buffer (BD Biosciences) according to the manufacturer’s instructions. Intracellular staining was performed with anti-α-smooth muscle actin (α-SMA)-eFluor570 (Invitrogen) and anti-cleaved caspase-3-Alexa Fluor 647 (Cell Signaling Technology) for 30 min at 4 °C, protected from light. Cells were washed twice in permeabilization buffer and resuspended in staining buffer prior to acquisition. Data were acquired on a Cytek Aurora, spectral flow cytometry (Cytek Biosciences) and analyzed using FlowJo software (BD Biosciences).

### Zymography assay

Matrix metalloproteinases (MMPs) activity in aortic tissue samples was assessed using the Novex™ Zymogram Plus Gelatin Gel Electrophoresis system (Thermo Fisher Scientific), following the manufacturer’s protocol. Briefly, equal amounts of protein extracted from aortic tissue homogenates were mixed with non-reducing sample buffer and loaded onto 10% gelatin-containing polyacrylamide gels. After electrophoresis, the gels were renatured and incubated in development buffer to allow enzymatic digestion of the gelatin substrate by active MMPs. Gels were then stained with Coomassie Brilliant Blue and destained to visualize zones of proteolysis, which appeared as clear bands against a blue background. Both the pro and active forms of MMP-2 and MMP-9 were identified based on their molecular weights. Band intensity was quantified using ImageJ software.

### Cytokine multiplex assay

Cytokine content in murine aortic tissue homogenates and culture supernatants were quantified using the Bio-Plex Bead Array system with a multiplex cytokine panel assay (Bio-Rad Laboratories, Hercules, CA), following the manufacturer’s instructions. Briefly, aortic tissues from various experimental groups were snap-frozen in liquid nitrogen, ground using a mortar and pestle, and homogenized in tissue extraction buffer. Protein concentration was determined using the BCA assay, and equal amounts of protein were used for cytokine measurements in the tissue samples. Similarly, macrophages were treated with 0.4 units/mL elastase for 5 minutes, washed twice with PBS, and cultured in serum-free medium for 24 hours with or without Protectin D1 (PD1; 300 nM). The culture medium was then collected and stored for subsequent cytokine analysis.

### Statistical analysis

Statistical analyses were performed using GraphPad Prism 10 (GraphPad Software, La Jolla, CA) software. For comparisons involving three or more groups, a one-way ANOVA followed by Tukey’s multiple comparisons test was utilized. Pairwise comparisons were conducted using either an unpaired Student’s t-test or, a non-parametric Mann-Whitney test. Data are presented as mean ± standard error of the mean, and statistical significance was defined as p < 0.05.

## RESULTS

### Single-cell RNA-sequencing analysis reveals dysregulation of PD1/GPR37 signaling-related genes in AAA patients

To evaluate evidence of PD1 and GPR37 receptor mediated signaling in human AAAs, single-cell RNA sequencing data from aortic tissue from AAA patients and comparative controls was analyzed using a previously reported sequencing dataset (GSE166676) (13), and differentially expressed genes were identified (Fig. 1A). Data was analyzed via Seurat and the macrophage clusters were identified for further analyses (Fig 1B). A total of 1530 genes were detected within the macrophage cluster and evaluated for differential expression. Of the 1530 genes, 610 were downregulated, 552 were upregulated, and 368 were unchanged (Supplemental Table S1). 7/610 downregulated genes and 9/552 upregulated were Related to Protectin D1 and GPR37 (Fig. 1D and 1E). Notably, this analysis suggests that genes known to regulate inflammation in AAAs (IFNG, CCR1, IL10, CCL2, P2RY6, CMKLR1, C5aR1, and CCR7) were not only dysregulated but also previously linked with PD1/GPR37-mediated signaling (9, 19, 20) These findings in human AAA tissue indicate the association between PD1/GPR37-mediated signaling in aortic inflammation, vascular remodeling and cell death and supported deciphering this pathway’s role in preclinical models of AAA.

**Figure 1:**
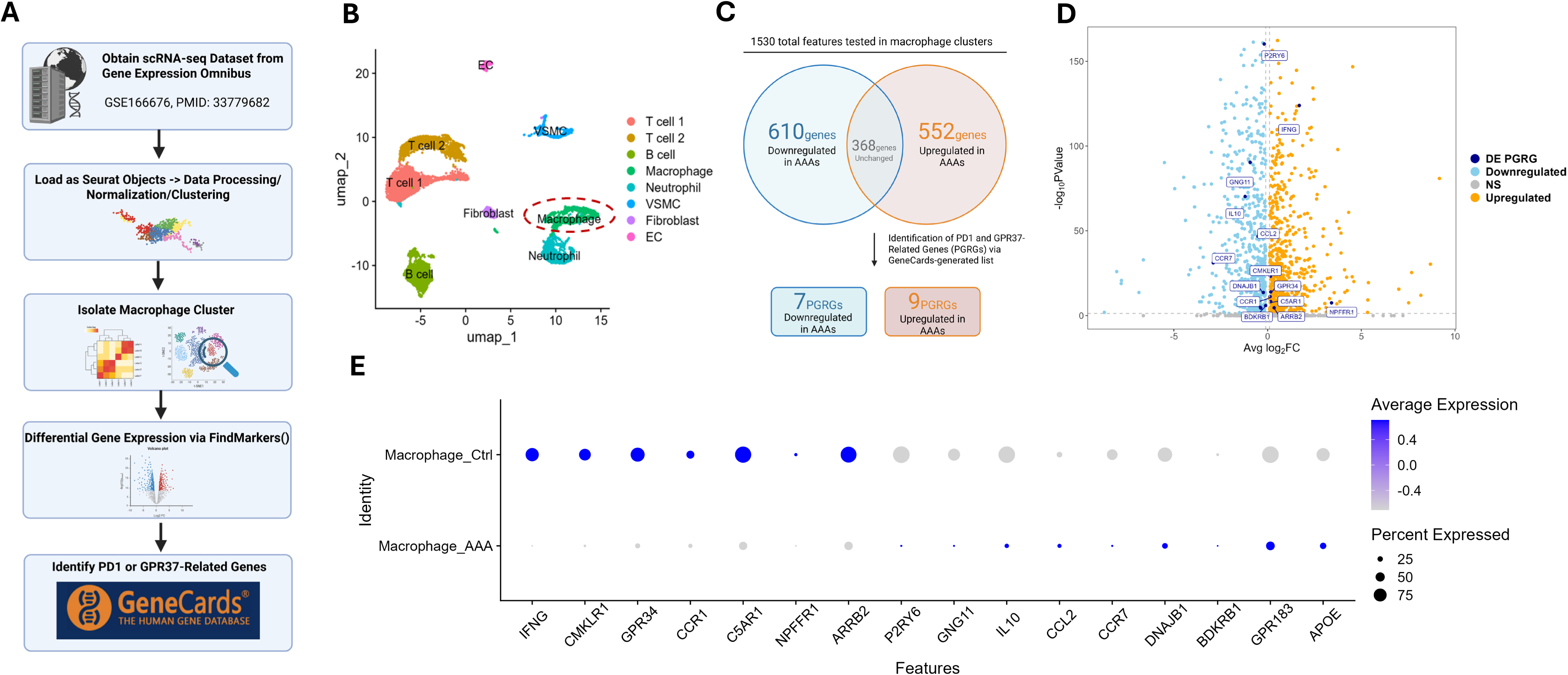
PD1 and GPR37 Receptor-related genes are dysregulated in macrophages of human AAAs. **A**. Bioinformatic workflow to re-analyze a publicly available scRNA-seq data set (GSE166676) from Gene Expression Omnibus. Data was analyzed via Seurat and the macrophage cluster was isolated. Differential expression analysis comparing macrophages from human AAAs (n=4) vs control aortas (n=2) was performed via FindMarkers calculated using Wilcoxon Rank Sum Testing. Genes with adjusted p-value < 0.05 and |fold-change| > 0.1 were considered significant (DEGs). A list of PD1 and GPR37 Receptor-related genes (PGRGs) were generated using GeneCards and used to identify differentially expressed PGRGs (DE PGRGs). **B.** Uniform manifold projection (UMAP) plot of the annotated clusters from human AAAs and control. The macrophage cluster is circled and subsetted for further analysis. **C.** Venn diagram displaying gene expression from this analysis. A total of 1530 genes were detected within the macrophage cluster and evaluated for differential expression. **D.** Volcano plot displaying significantly downregulated (skyblue) and upregulated (orange) genes. DE PGRGs are highlighted and colored (navy blue). **E.** DotPlot of all 30 DE PGRGs in macrophages displayed as average expression aggregated between conditions (Ctrl vs AAA).

### PD1 treatment mitigates aortic inflammation and vascular remodeling during AAA formation

Using the topical elastase AAA model, wild-type (WT) male mice were treated with either elastase or heat-inactivated elastase on day 0 and injected with vehicle or PD1 (300 ng/mouse/day) on postoperative days 1,3,5,7 and 9 and harvested on day14 (Fig. 2A). A significant increase in mean aortic diameter was noted in elastase-treated mice compared to heat-inactivated elastase-treated control mice (156±10.3 vs. 4.4±1.5%; Fig. 2B-C). Mice treated with PD1 demonstrated significantly decreased mean aortic dilation compared to mice treated with vehicle alone (117±7.9 vs. 156±10.3%; Fig. 2B).

**Figure 2.**
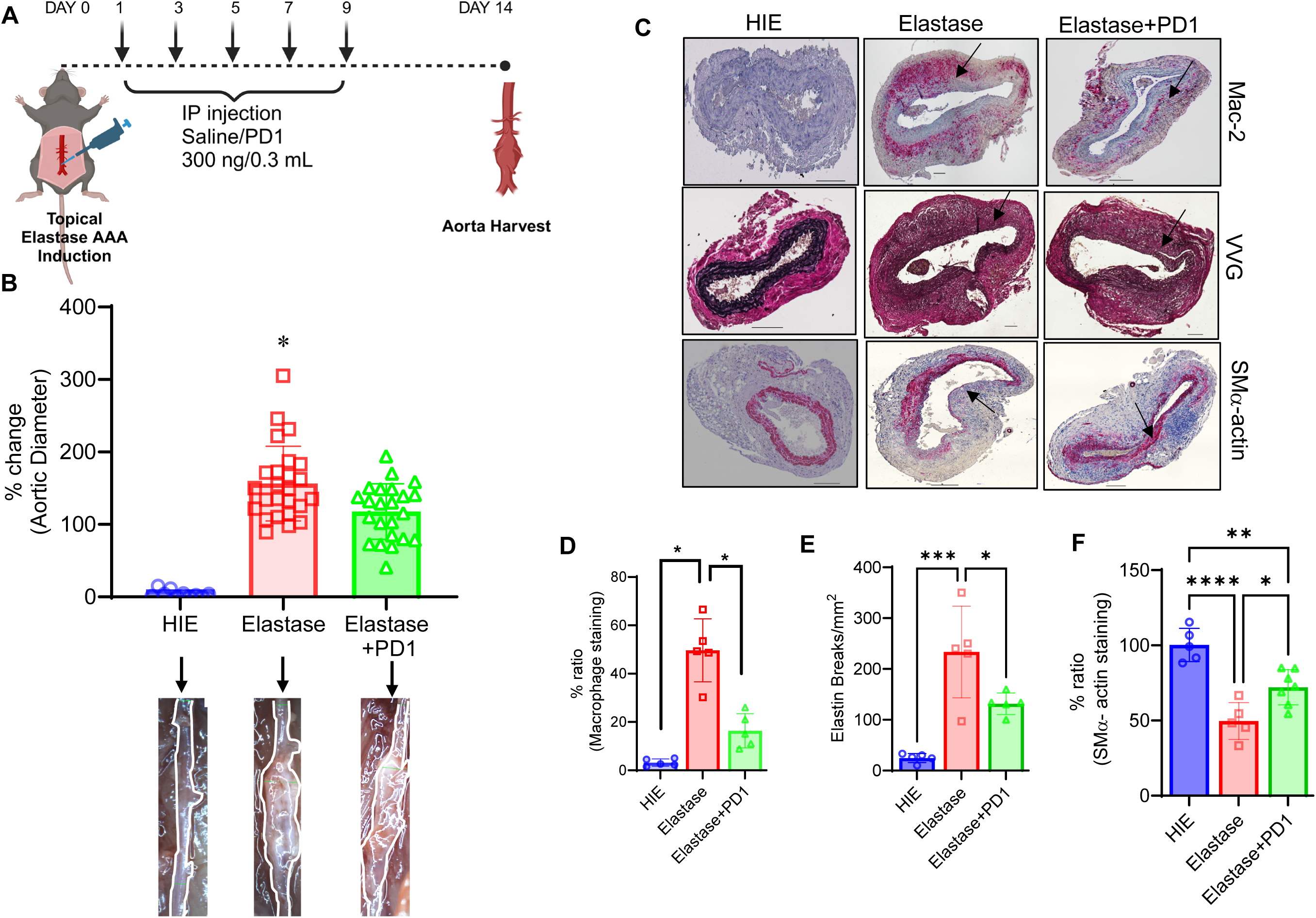
PD1 administration mitigates AAA formation. **A.** Schematic representation of the elastase-treatment AAA model. WT mice were treated with either elastase or heat-inactivated elastase (HIE). **B.** Elastase-treated mice exhibited significant increase in aortic dilation compared to controls, that was significantly attenuated by PD1 administration. Representative images of aortas from each group are shown. n=10-25/group; *p<0.05 vs. other groups. **C-D.** Histological analysis and quantification of aortic sections on day 14 demonstrating increased macrophage infiltration (Mac-2 staining), disrupted elastic fibers (Verhoeff–Van Gieson staining), and reduced smooth muscle cell α-actin (SM α-actin) expression in elastase-treated aortas was significantly ameliorated after PD1 treatment. n=5/group; *p<0.02; **p=0.002; ***p=0.001; ****p<0.0001. HIE, heat-inactivated elastase.

Furthermore, PD1 treatment led to a significant increase in smooth muscle alpha-actin (SMα-actin) expression (72.1 ± 11.6 vs. 49.7 ± 12.2%), a reduction in elastin fragmentation (131 ± 21 vs. 233 ± 90 number of breaks/mm^2^), and decreased macrophage infiltration (16 ± 9 vs. 49 ± 13%) on day 14 compared to vehicle-treated mice (Fig. 1C-D). Moreover, the aortic tissue expression of pro-inflammatory cytokines (IL-1β, TNF-α, MCP-1, MIP-2, IL-17, IL-6) and matrix metalloproteinases, MMP2 but not MMP9, were significantly attenuated by PD1 treatment (Fig. 3A-C and (Supplementary Fig. S1). Mechanistically, elastase-treated aortic tissue demonstrated a significant upregulation of SMC-dependent apoptosis compared to controls, and PD1 treatment significantly enhanced the clearance of apoptotic SMCs by macrophages via efferocytosis in WT mice compared to untreated controls (Fig. 3D-E and Supplementary Fig. S2-S3).

**Figure 3.**
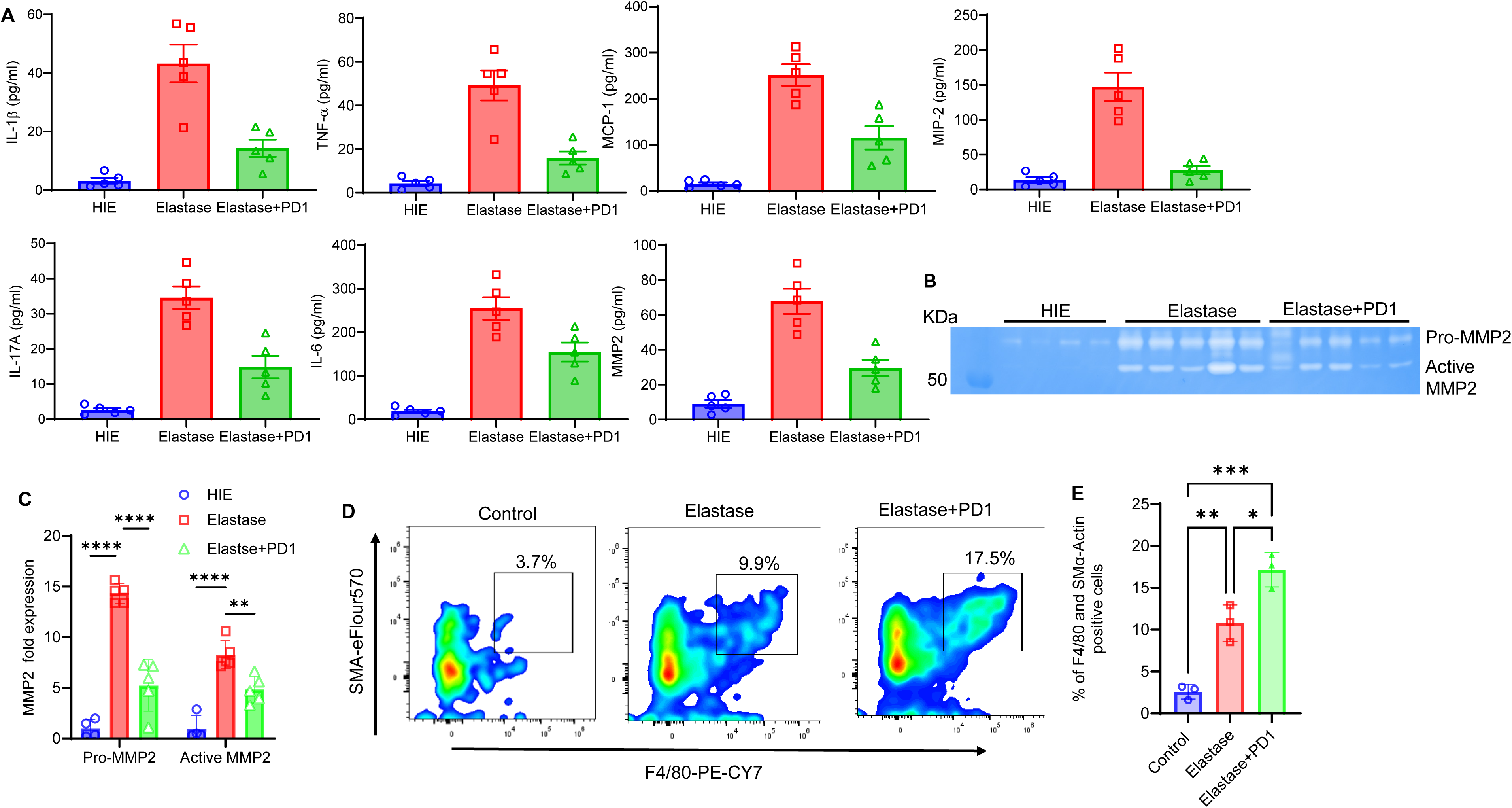
PD1 administration ameliorates proinflammatory cytokine/chemokine and MMP production. **A**. Pro-inflammatory cytokine and MMP2 expressions in aortic tissue of WT mice treated with PD1 were significantly attenuated compared to elastase treatment alone. *p<0.01 vs. other groups; n=5/group. **B-C.** MMP-2 expression levels in aortic tissues were assessed by gelatin zymography. A significant reduction in active MMP-2 was observed after PD1 treatment compared to untreated group. ****p<0.01; **p<0.05; n=5/group. **D-E.** Flow cytometry analysis of aortic tissue demonstrates enhanced efferocytosis and uptake of apoptotic SMCs by macrophages after PD1 treatment compared to untreated controls. *p<0.05; **p<0.03; ***p<0.001; n=3/group.

### PD1 mediated protection of AAA is regulated by GPR37 receptors

To determine whether GPR37 receptors mediate the protective effects of PD1 in AAA, we utilized GPR37⁻^/^⁻ mice in the elastase-treatment AAA model. In contrast to WT mice, PD1 administration did not result in a significant reduction in aortic diameter in GPR37⁻^/^⁻ mice compared to vehicle controls (121.3±37.6 vs 110.3±21.7%; Fig 4A). Histological analyses further revealed no significant differences in SMα-actin (22.9±7.4 vs 20±3.7%), elastin breaks (194±62 vs 236±49 number of breaks/mm^2^) and macrophage infiltration (49.9±9.2 vs 41.6±11.1%) between PD1- and vehicle-treated GPR37⁻/⁻ mice (Fig. 4B-E).

**Figure 4.**
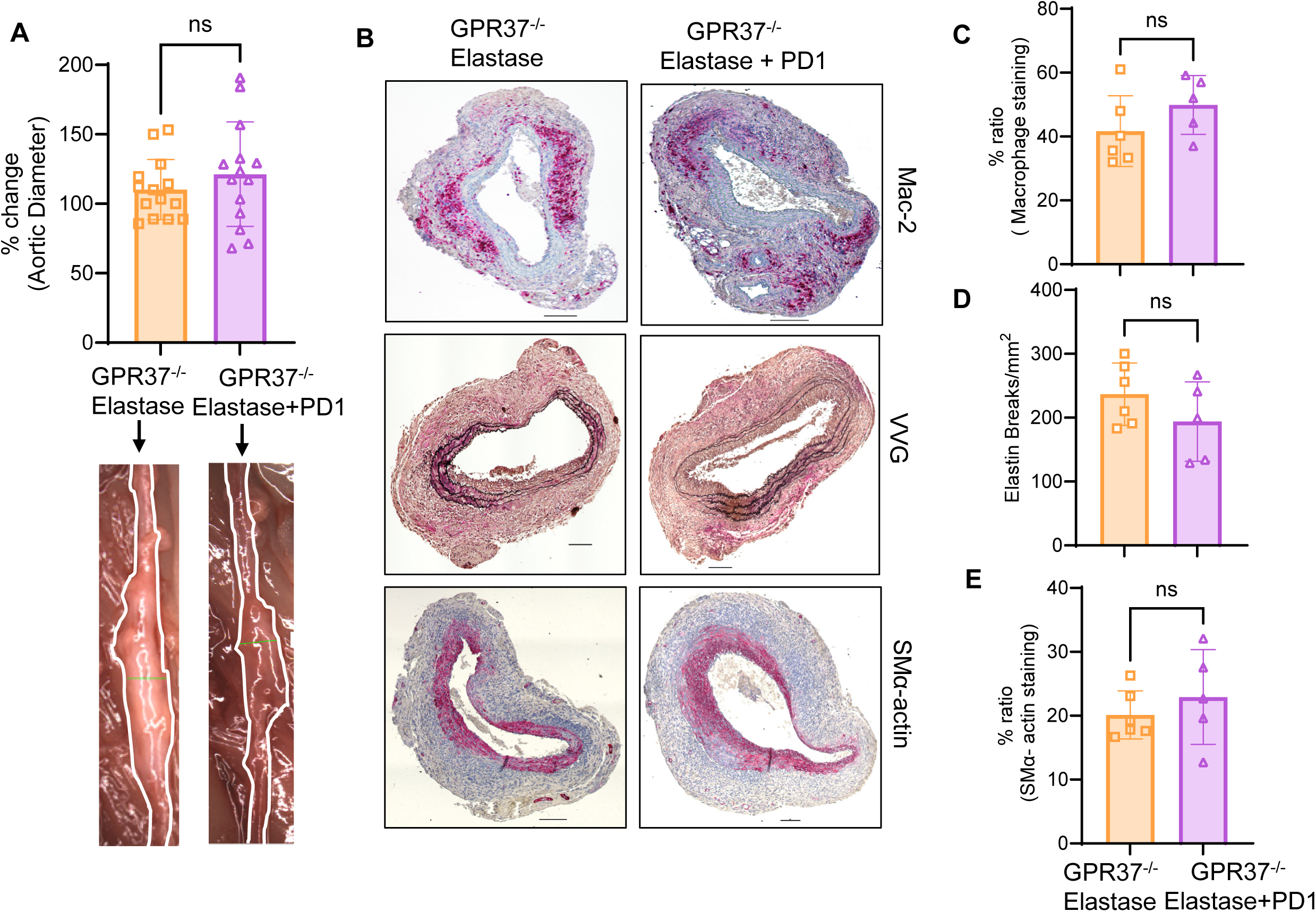
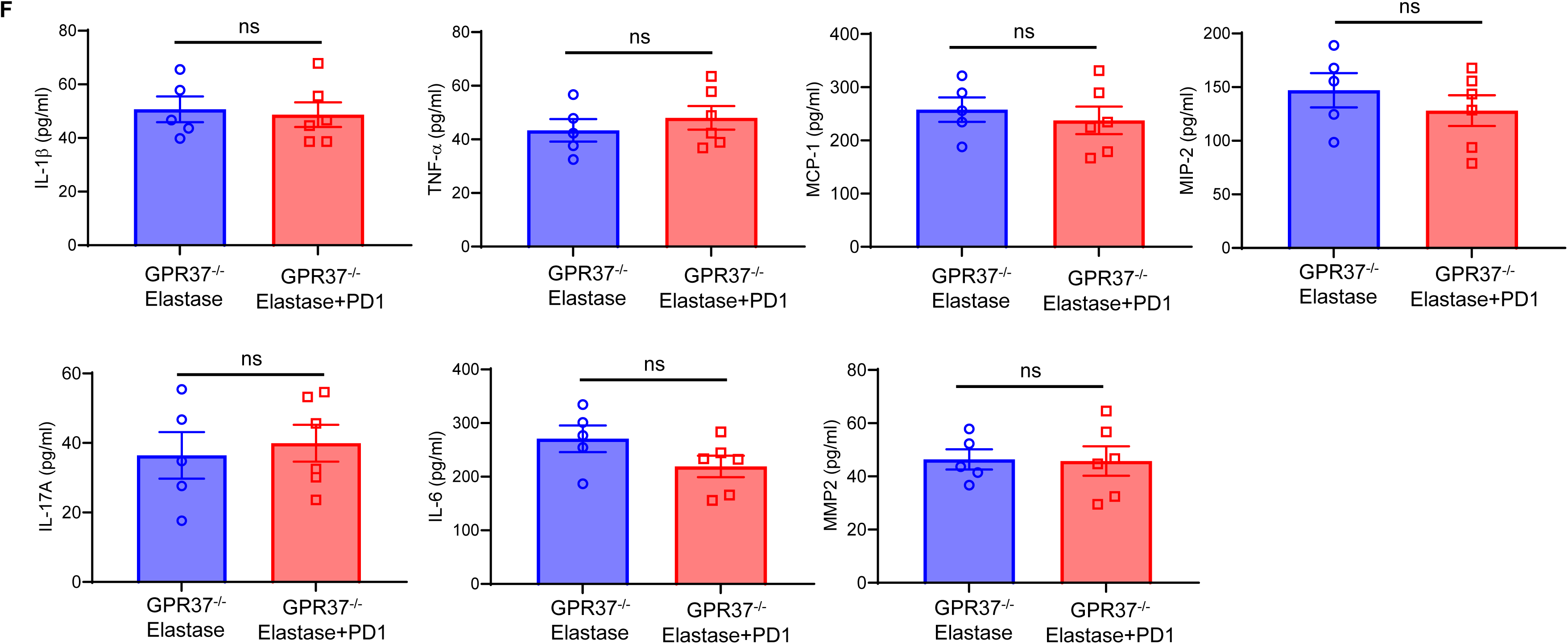
PD1-mediated protection against AAA is dependent on GPR37 receptors. **A.** GPR37⁻^/^⁻ mice were subjected to elastase-induced AAA and treated with or without PD1. Aortic tissues were harvested on day 14 for analysis. No significant changes in aortic dilation were observed after PD1 treatment compared to untreated controls. Representative images of aortas from each group are shown. **B.** In PD1-treated GPR37^-/-^ mice, comparative histology demonstrated no differences in SMα-actin expression, elastin fiber disruption, and immune cell infiltration macrophage staining) compared to untreated mice. Arrows point to areas of immunostaining. Scale bar=100μM. **C-E.** Quantification of immunohistochemical staining demonstrates no significant differences between SMα-actin staining, elastin breaks, and macrophage staining in PD1-treated aortic tissue compared to mice treated with elastase alone. ns, not significant; n=5/group. **F.** Pro-inflammatory cytokine and MMP2 expressions in aortic tissue of GPR37^-/-^ mice treated with PD1 demonstrated no differences compared to elastase treatment alone. ns, not significant; n=5/group.

Additionally, no significant change in the expression of proinflammatory cytokines and MMP2 in aortic tissue of PD1-treated GPR37^-/-^ mice was observed compared to elastase-treated GPR37^-/-^ mice alone (Fig. 4F). Gelatin zymography of aortic tissues demonstrated that MMP2 and MMP9 activity levels remained unchanged with PD1 administration in GPR37⁻/⁻ mice, further supporting the lack of PD1-mediated protection in the absence of GPR37 (Supplementary Fig. S4). These findings collectively indicate that the protective effects of PD1 in AAA are largely mediated through GPR37 signaling, and that PD1 fails to confer vascular protection in the absence of this receptor.

### PD1 confers protection against preformed AAAs

Given the significant reduction in aortic dilation observed with PD1 administration in the acute AAA model, we next investigated whether PD1 could confer protection against already established aneurysms using a chronic AAA model. In this model, PD1 (300 ng/mouse/day) was administered starting on day 14 post-surgery and continued daily until day 28 and aortic tissue was harvested on day 28 for analysis (Fig. 5A). PD1 treatment resulted in a significant reduction in aortic dilation compared to saline-treated controls (261±133 vs. 470±134, Fig. 5B). Histological analysis and quantitative assessments revealed that PD1-treated mice exhibited increased expression of SM-α actin, as well as reduced elastin fragmentation and macrophage infiltration, indicating enhanced vessel integrity (Fig. 5C-F). Furthermore, a significant attenuation of pro-inflammatory cytokine and MMP2 expressions was observed in aortic tissue of PD1 treated mice compared to untreated controls (Fig. 5G). Collectively, these findings indicate that PD1 provides vascular protection not only during the early phases of AAA development but also in established aneurysms, potentially through preservation of smooth muscle integrity, reduced inflammation, and modulation of matrix-degrading enzymes.

**Figure 5.**
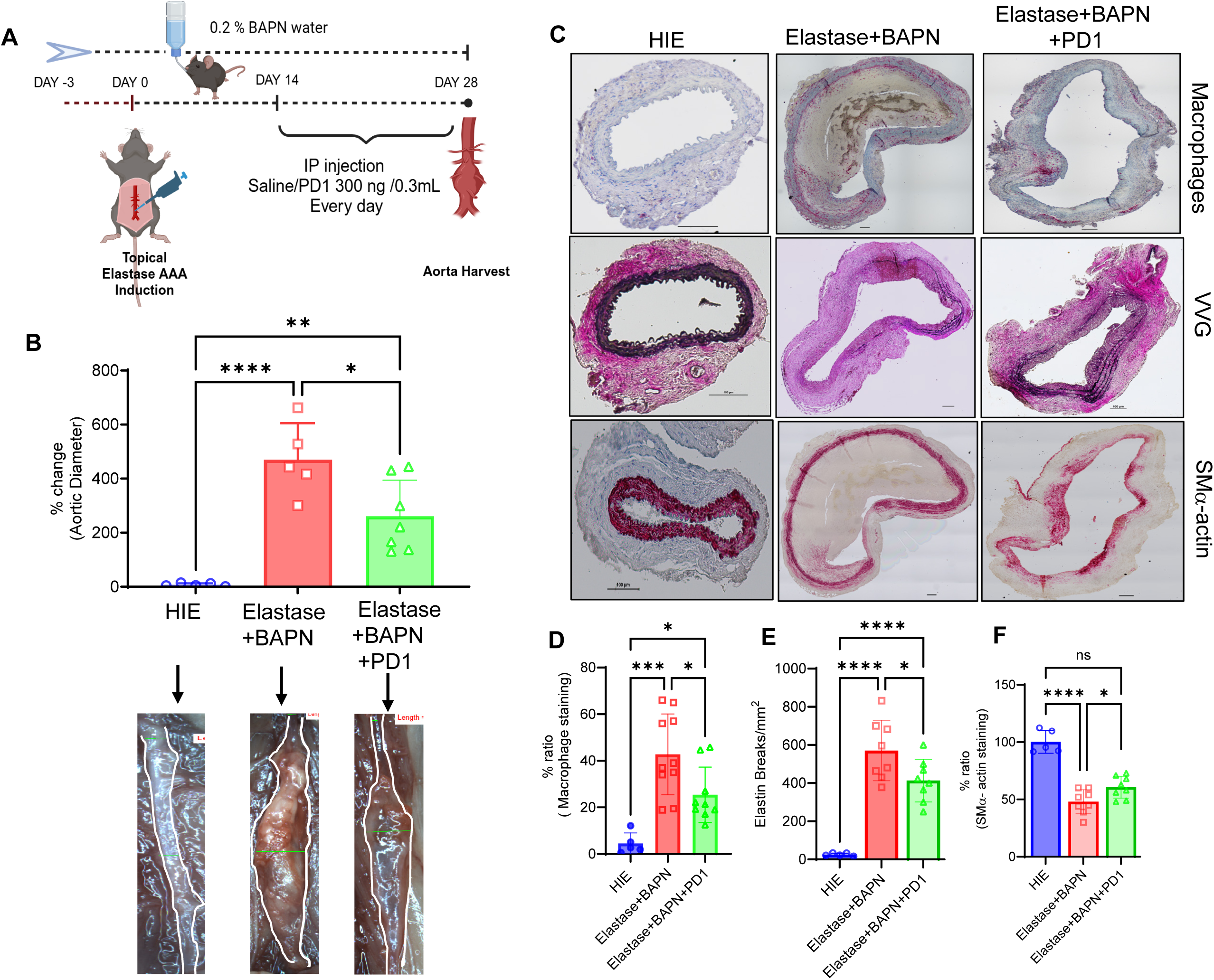
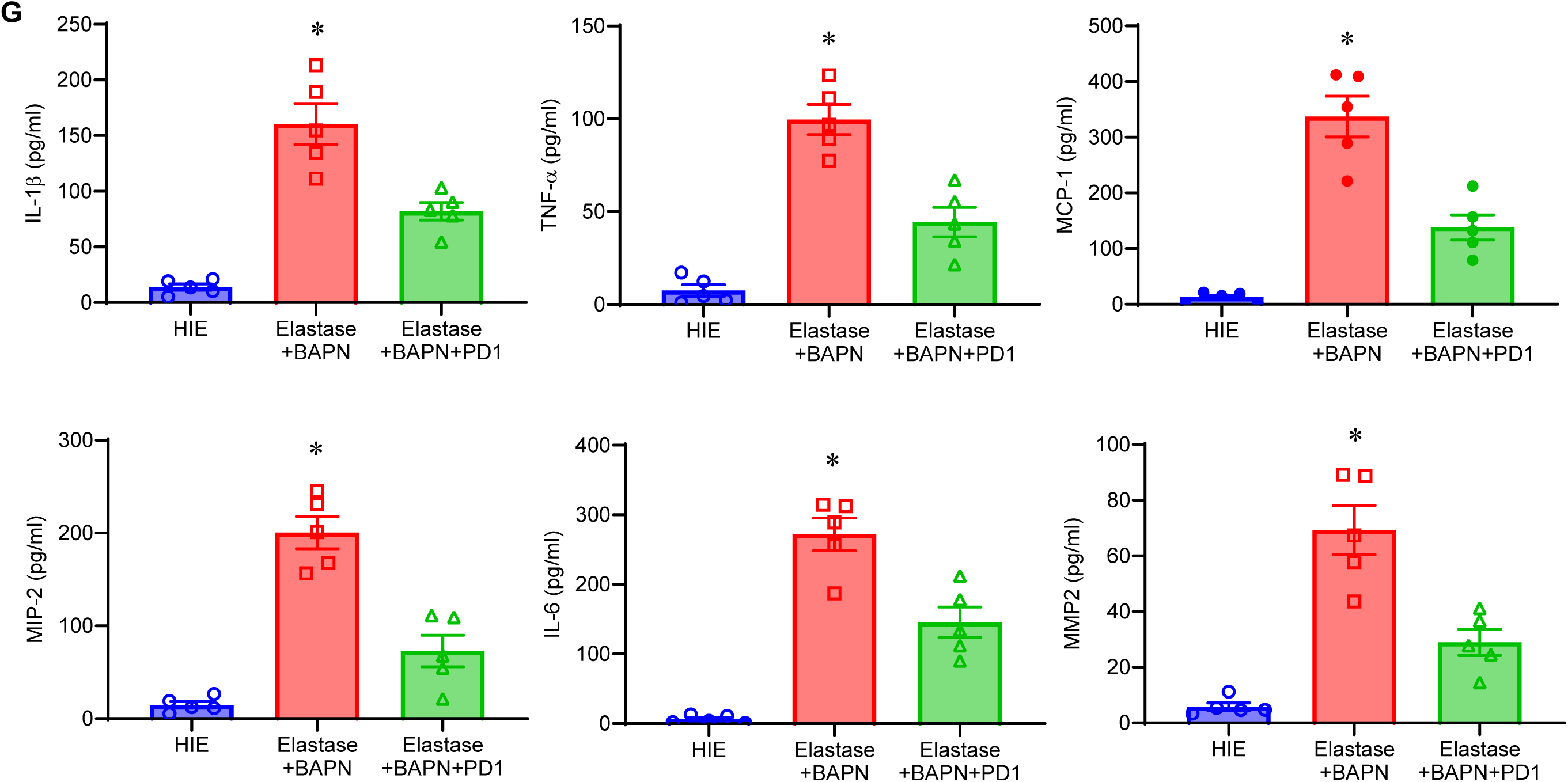
PD1 administration mitigates pre-formed AAAs. **A.** Schematic of the BAPN + elastase-induced chronic AAA model. **B.** Aortic diameter is significantly attenuated in PD1-treated mice compared to elastase+BAPN treated mice on day 28. *p=0.01; **p=0.004; ****p<0.0001; n=5/group. Comparative representative of aortic phenotype in each group is shown. **C.** PD1 treatment preserves aortic morphology in the chronic murine AAA model. In PD1-treated mice, comparative histology demonstrated increased SMα-actin expression, decreased elastin fiber disruption, and reduced immune cell infiltration (macrophage staining) compared to untreated mice. Arrows indicate areas of immunostaining. **D-F.** Quantification of immunohistochemical staining demonstrates increased SMα-actin staining, and reduced elastin breaks, and macrophage staining in PD1-treated tissue compared to mice treated with elastase+BAPN alone. Scale bar=100μM.*p=0.03; ***p<0.0001; ****p<0.0001; ns, not significant; n=5/group. **G.** Aortic tissue from PD1-administered WT mice in the elastase+BAPN-treated group showed significant decrease in pro-inflammatory cytokine/chemokine production and MMP2 expressions compared to elastase+BAPN-treated WT mice. *p<0.05 vs. other groups, n=5/group,

### PD1 enhances macrophage-dependent efferocytosis via GPR37

To investigate the clearance of apoptotic SMCs by macrophages via efferocytosis, and its modulation by PD1, we performed *in vitro* efferocytosis assays using peritoneal macrophages isolated from wild-type (WT) and GPR37⁻^/^⁻ mice. Murine macrophages were used as phagocytes, and apoptotic MOVAS cells served as the target population. The two cell types were co-cultured in the presence or absence of PD1, and efferocytosis was assessed by measuring the percentage of macrophages (F4/80⁺) that engulfed PKH67-labeled apoptotic MOVAS cells (Fig. 6A and (Supplementary Fig. S5). In WT macrophages, PD1 treatment significantly enhanced efferocytic capacity compared to vehicle-treated controls (65.7 ± 8.4% vs. 51.5 ± 6.3%; Fig. 6B). However, in GPR37⁻^/^⁻ macrophages, PD1 treatment did not significantly alter efferocytosis rates relative to controls (29.0 ± 6.3% vs. 30.6 ± 4.7%, Fig. 6C). Moreover, PD1 treatment of macrophages attenuates pro-inflammatory cytokine production such as IL-1β and TNF-α and enhanced anti-inflammatory IL-10 expression in a GPR37-dependent manner (Fig. 6D-E and Supplementary Fig. S6). These findings suggest that PD1 enhances macrophage-mediated efferocytosis in a GPR37-dependent manner and attenuates pro-inflammatory paracrine secretion that mitigates SMC-dependent remodeling to protect against AAA formation and aortic rupture (Fig. 6E).

**Figure 6.**
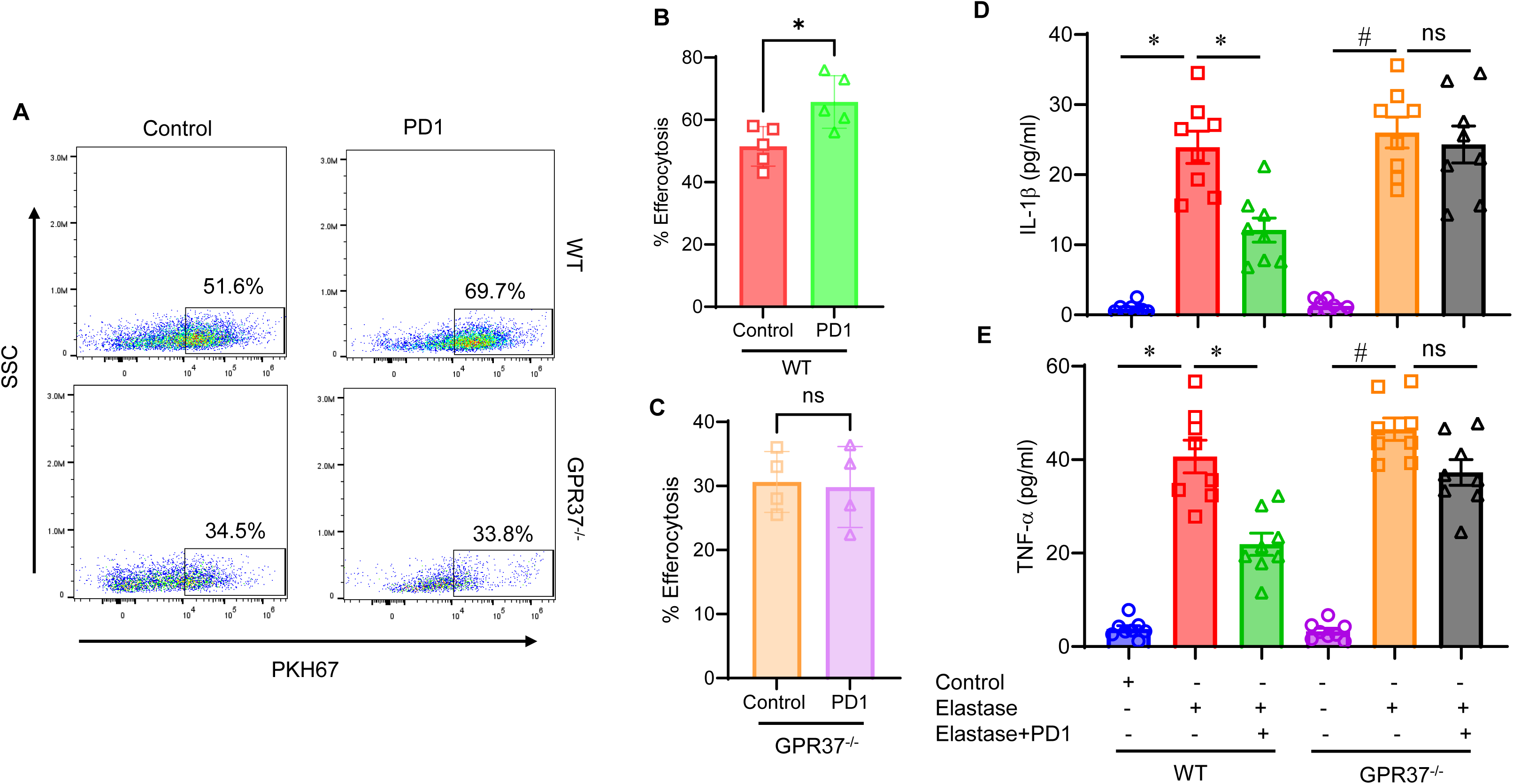
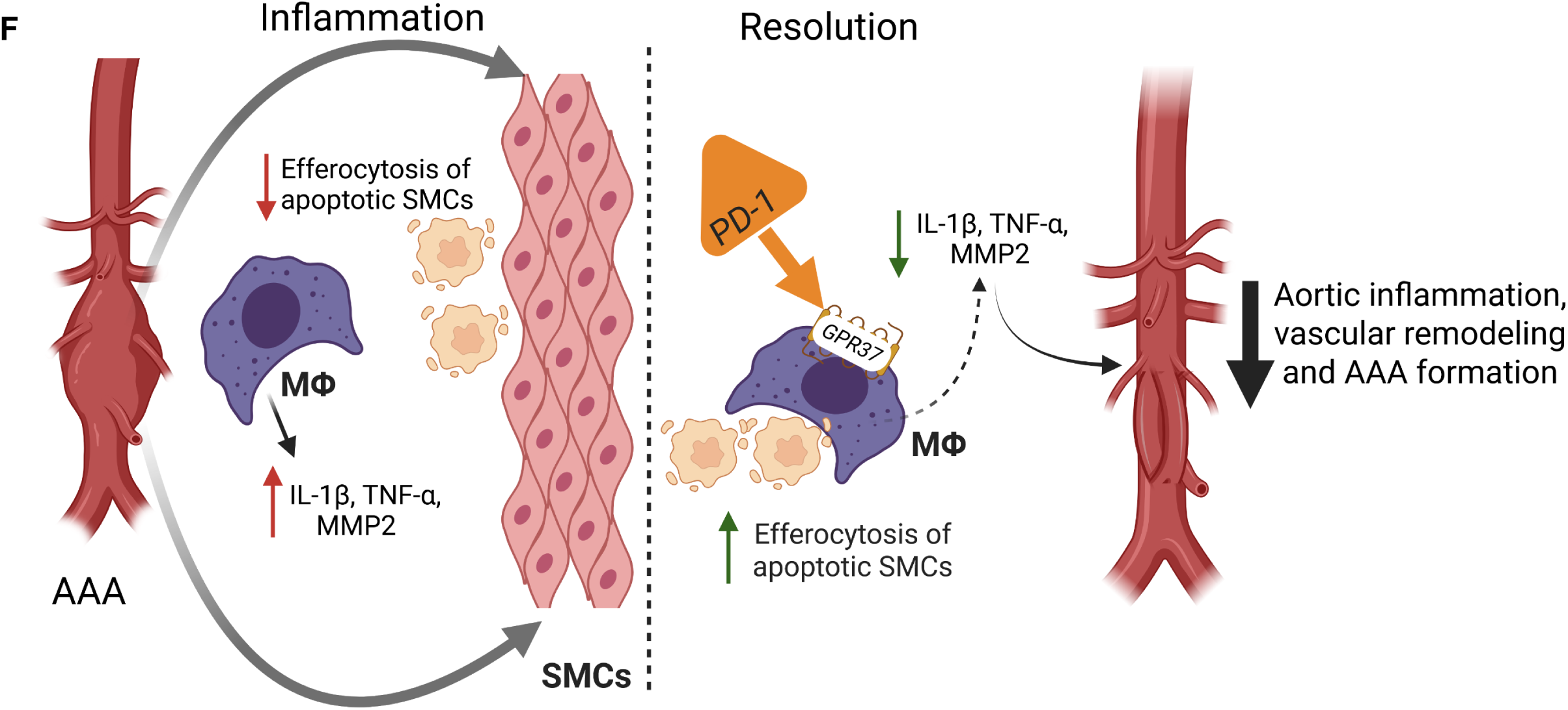
PD1 enhances macrophage-dependent efferocytosis in a GPR37-dependent manner. A-C. *In vitro* efferocytosis was performed using peritoneal macrophages isolated from WT and GPR37⁻^/^⁻ mice. Apoptotic SMCs stained with PKH67 were used as bait. PD1 treatment significantly enhanced efferocytosis in WT, but not GPR37^-/-^, macrophages compared to untreated controls. **D-E.** Purified macrophages from WT and GPR37^-/-^ mice were exposed to transient elastase-treatment and expression of IL-1β and TNF-α was significantly increased in culture supernatants compared to respective controls. PD1 treatment significantly attenuates cytokine expressions in WT but not GPR37^-/-^ macrophages. *p<0.03; #p<0.001; ns, not significant; n=8/group. **F.** Schematic description of PD1-mediated attenuation of AAA formation. The activation of macrophages leads to release of pro-inflammatory cytokines and dysregulation of efferocytosis of apoptotic SMCs that modulates vascular remodeling during AAA formation. During the resolution process, exogenous PD1 acts on GPR37 receptors on macrophages to upregulate efferocytosis of apoptotic SMCs during AAA formation. This process downregulates IL-1β and TNF-α secretion that leads to SMC preservation, decrease in MMP2 activation and subsequent aortic remodeling as well as AAA formation. PD1, protectin D1; GPR37, G-protein coupled receptor 37; SMCs; smooth muscle cells; MФ; macrophages; MMP; matrix metalloproteinases.

## DISCUSSION

This study demonstrates that the process of inflammation-resolution can be mediated by SPMs, such as PD1, via GPR37 receptor-dependent signaling that involves upregulating macrophage-dependent efferocytosis of apoptotic SMCs to mitigate aortic inflammation, vascular remodeling and AAA formation. Single cell RNA analysis of human AAA tissue demonstrated dysregulation of several PD1/GPR37 associated genes providing association between endogenous inflammation and resolution pathways. Importantly, *in vivo* and *in vitro* studies demonstrate the ability of PD1 to pharmacologically modulate macrophage-mediated efferocytosis and downregulate proinflammatory cytokine paracrine secretion that decreases MMP2 expression and vascular remodeling. Collectively, these results suggest an important role of efferocytosis in AAA and the ability of PD1/GPR37 signaling to mitigate AAA formation and prevent aortic rupture.

Dysregulated resolution of inflammation can trigger a cascade of pathological events, including persistent tissue damage and impaired clearance of apoptotic cells, ultimately contributing to chronic inflammatory cardiovascular disorders (21, 22). Once considered a passive process, inflammation resolution is now recognized as an active, highly coordinated sequence of events driven by SPMs, such as lipoxins, resolvins, maresins, and protectins (23). SPMs exert potent pro-resolving and anti-inflammatory effects, in part by enhancing macrophage-mediated clearance of apoptotic cells through efferocytosis to maintain tissue homeostasis (24). For example, we have previously demonstrated that Resolvin D1 (RvD1)-treatment can mitigate aortic inflammation, decrease immune cell infiltration, reduce MMP activity, and promote macrophage polarization toward an anti-inflammatory phenotype (25). Additionally, our studies demonstrated that RvD1 decreased AAA formation through inhibition of neutrophil extracellular traps (NETosis) which is a downstream event of IL-1β mediated neutrophil activation and infiltration in the aortic wall (26, 27). SPMs exert their protective effects mediated via specific receptors that elucidate anti-inflammatory activities in a cell- and tissue-specific manner (28–32). RvD1 has been shown to exhibit its pro-resolving function by signaling through two G protein-coupled receptors (GPCRs), A lipoxin/formyl peptide receptor 2 (ALX/FPR2) and GPR32 that have been shown to decrease PMN infiltration as well as stimulating macrophage phagocytic function (33). Furthermore, our recent findings provide evidence for MaR1-mediated ingestion of apoptotic SMCs via LGR6 activated macrophages that stimulate TGF-β signaling and protect SMC activation (34). These studies implicate the effects of disrupted SMC TGF-β signaling that can be enhanced by SPMs to mediate resolution of aneurysm formation via SMC reprogramming.

The current study investigated the mechanistic role of PD1 and its downstream signaling via GPR37 in AAA progression and resolution failure. Although GPR37 has traditionally been classified as an orphan receptor, emerging evidence reveals its interaction with PD1 in macrophages, suggesting a previously unrecognized role in inflammation resolution.(10, 35). Recent studies have reported that PD1 activation of GPR37 enhances macrophage phagocytic capacity, facilitates pain resolution, and improves outcomes in sepsis models (9, 10). In the context of AAA, inflammation-driven degradation of the aortic wall is exacerbated by oxidative stress, vascular VSMC apoptosis, and extracellular matrix remodeling. These processes promote the expression of ECM-degrading enzymes, particularly MMP2 and MMP9-produced by VSMCs and infiltrating macrophages, respectively (36).Our results show that PD1 treatment significantly reduces aortic inflammation, leukocyte infiltration, and pathological vascular remodeling during AAA formation in a GPR37-dependent manner. Importantly, the macrophage-mediated clearance of apoptotic cells, an essential feature of inflammation resolution, is suppressed in AAA but is restored by PD1/GPR37 signaling. PD1 exhibits robust therapeutic potential in AAA by targeting multiple pathological mechanisms, including dampening of inflammation, preservation of vascular integrity, and enhancement of apoptotic cell clearance. These structural benefits are associated with reduced expression of pro-inflammatory cytokines and matrix metalloproteinases (MMPs), consistent with the known actions of SPMs (37).

Macrophages demonstrate varied phenotypes, such as proinflammatory M1-like phenotypes and anti-inflammatory M2 phenotypes, as well as many intermediate phenotypes that mediate the detrimental and protective actions of MΦs (15, 16). Our study has shown a critical role of GPR37 in regulating macrophage phenotypes via GPR37 to demonstrate resolution of M2-like phenotypes that express decreased paracrine secretions such as IL-1β and TNF-α, but increased levels of IL-10. A critical component of this switch is efferocytosis, which reprograms macrophages to release pro-resolving mediators (38). GPR37 has recently been shown to be expressed by CD163⁺ macrophages in the dorsal root ganglia neurovascular unit, contributing to pain resolution (39). These findings identify GPR37 as a critical mediator of PD1-dependent protective actions and reinforce its emerging role in macrophage function and tissue repair (9). Given the central role of SMC apoptosis in aneurysm pathophysiology and its association with unresolved inflammation and defective efferocytosis (40), PD1 enhanced apoptotic cell clearance by macrophages in a GPR37-dependent manner, thereby implicating this axis in the macrophage-parenchymal cell crosstalk during AAA formation. This selective response reveals a novel immunoregulatory function for GPR37 in resolution of AAAs as well as protection against aortic rupture (10).

It is important to note that a few limitations exist in this study. The modulation of endogenous inflammation-resolution in human AAAs can be regulated by the balance of several lipid species that are affected due to the inflammatory milieu as well as underlying comorbidities.

Therefore, using bioactive lipids as an exogenous pharmacotherapy may likely involve a multitude of proresolving bioactive lipids as well as sequential administration over a chronic duration to circumvent the leukocyte-parenchymal interactions in chronic inflammation. Accordingly, clinical translation to human subjects remains to be determined in a large preclinical animal model to determine the safety and efficacy of PD1-mediated immunomodulation. Furthermore, a combined therapeutic strategy using various bioactive isoforms of SPMs represents an effective strategy to treat and mitigate aortic inflammation and vascular remodeling in chronic vascular diseases like AAA. Our recent findings and the present study indicate that the comparative efficacy of specific subsets of omega 3-fatty acid derived bioactive lipids can act on different immune and parenchymal cells. Therefore, a synergistic approach of RvD1, MaR1 and PD1 on aortic inflammation as a multifaceted approach for a combined therapeutic intervention should be delineated in a sex-dependent large animal model as a follow up study to decipher a clinically translatable strategy.

Collectively, our findings establish PD1 as a potent modulator of AAA pathobiology and identify GPR37 as an essential receptor mediating its therapeutic effects. By targeting inflammation and defective efferocytosis, PD1 holds significant promise for both the prevention and treatment of AAA. The mechanistic approach of PD1-mediated mitigation of aortic inflammation is conferred through increased efferocytosis of apoptotic vascular SMCs within the aortic wall via PD1-GPR37 interaction providing a mechanistic and non-surgical approach for preventing aortic ruptures. Further investigation is pivotal to demonstrate the combined therapy of specific SPM bioactive isoforms that are likely to provide insight for a clinically translatable approach in aortic aneurysms.

## Acknowledgments

The authors would like to thank Tabitha Randi for her impeccable help with administrative duties and laboratory management.

## Supplementary Material

Supplementary material (Figures S1-S6 and Table S1) are available at PNAS Nexus online.

## Author contributions

A.A., M.F., G.L., J.A.V., G.S., S.S., W.U. and A.K.S. conducted the research, performed experiments, collected, analyzed and interpreted data. A.A., A.K.S., and G.R.U. were instrumental in writing and editing the manuscript. A.K.S. and G.R.U. conceived and designed the study, supervised the project, obtained research funding and have access to all the data. All authors read the final manuscript and gave final approval of the version submitted.

## Data Availability

All relevant data are included in the manuscript or the Supplementary material.

