## Supplemental Figures S1-S6 for "Protectin D1/GPR37 signaling enhances macrophage-dependent efferocytosis to attenuate experimental abdominal aortic aneurysm formation"

**Supplementary Figure S1**

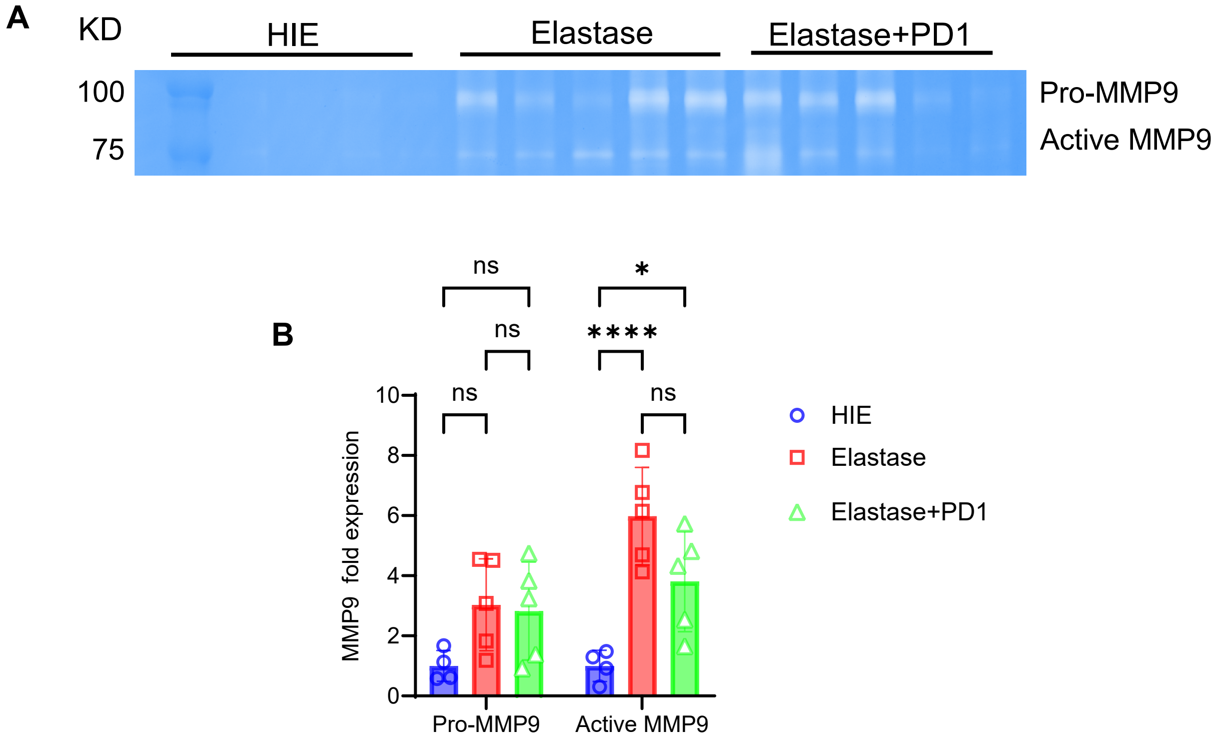


**Supplementary Figure S1. A.** Gelatin zymography of murine aortic tissue in WT mice and **B,** quantification of optical density (O.D.) shows no significant change in levels of proMMP9, but a significant increase in active MMP9 expression, compared to controls, that was not altered by PD1 treatment on day 14. n=4-5/group; *p<0.01; ****p<0.001; ns, not significant; HIE, heat-inactivated elastase.

### Supplementary Figure S2

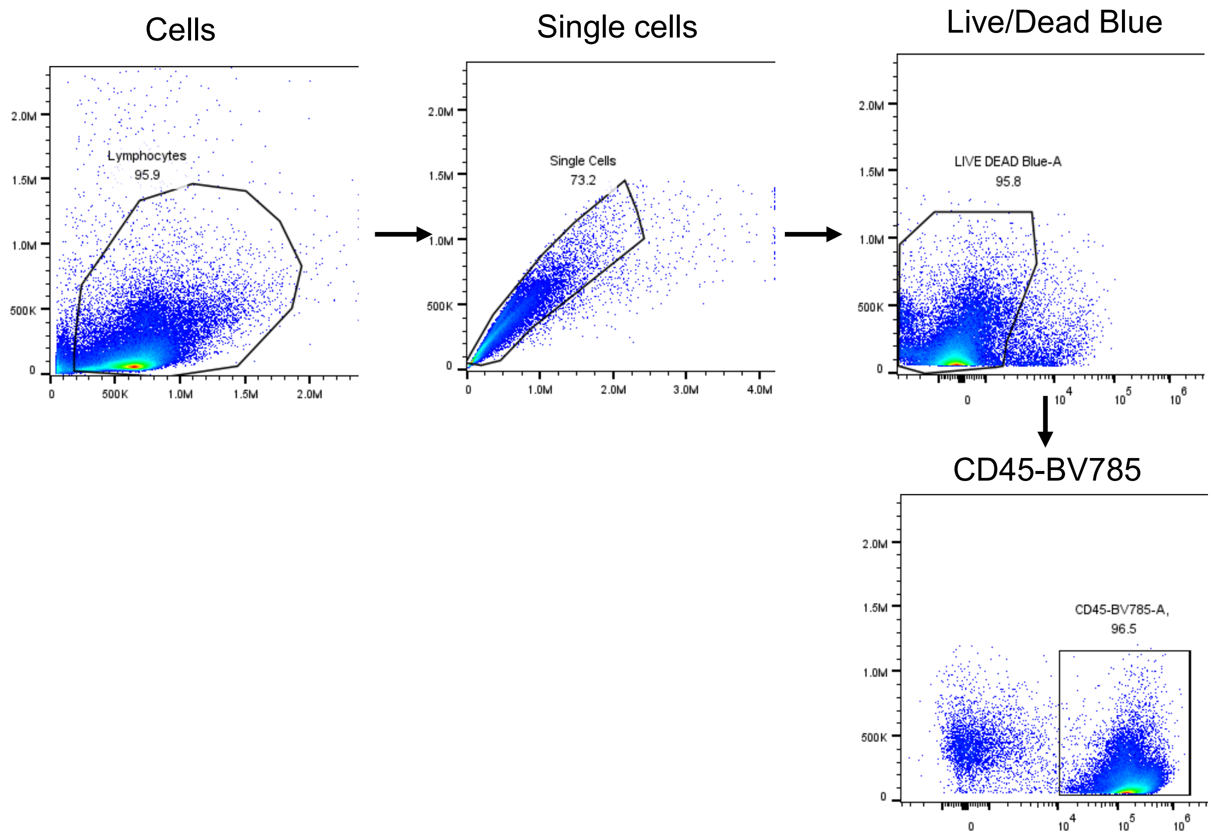


**Supplementary Fig. S2**. Gating strategy for flow cytometry analysis of efferocytosis using *in vivo* aortic tissue.

### Supplementary Figure S3

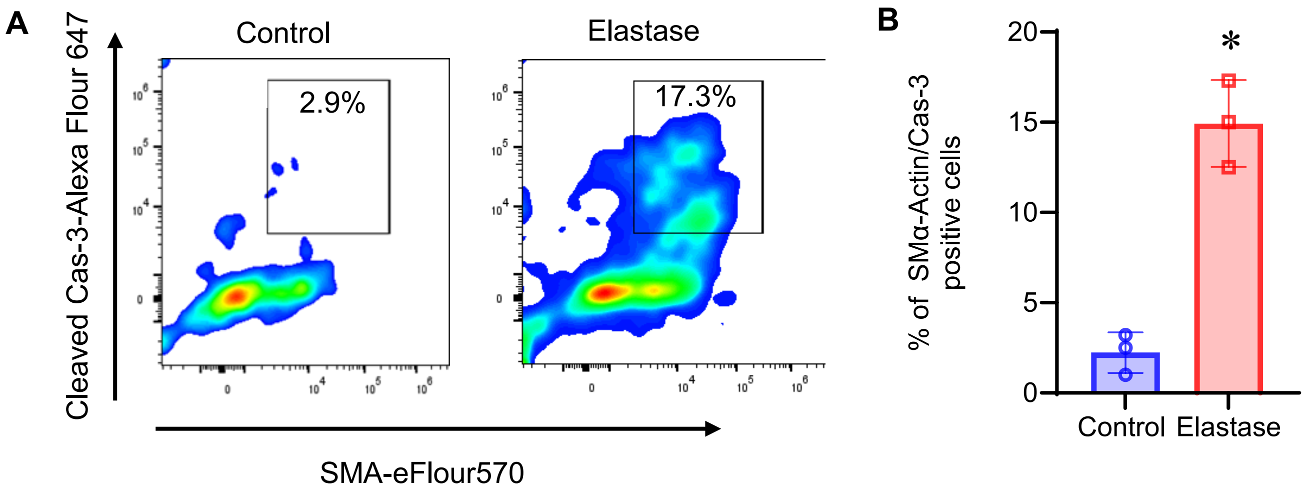


**Supplementary Fig. S3**. **A.** Flow cytometry analysis showing a significant increase in cleaved caspase-3 expression in SMCs following elastase-treatment compared to controls in aortic tissue on day 3. **B.** Quantification of apoptotic SMCs in aortic tissue of elastase-treated WT mice compared to controls. *p<0.01; n=3/group.

### Supplementary Figure S4

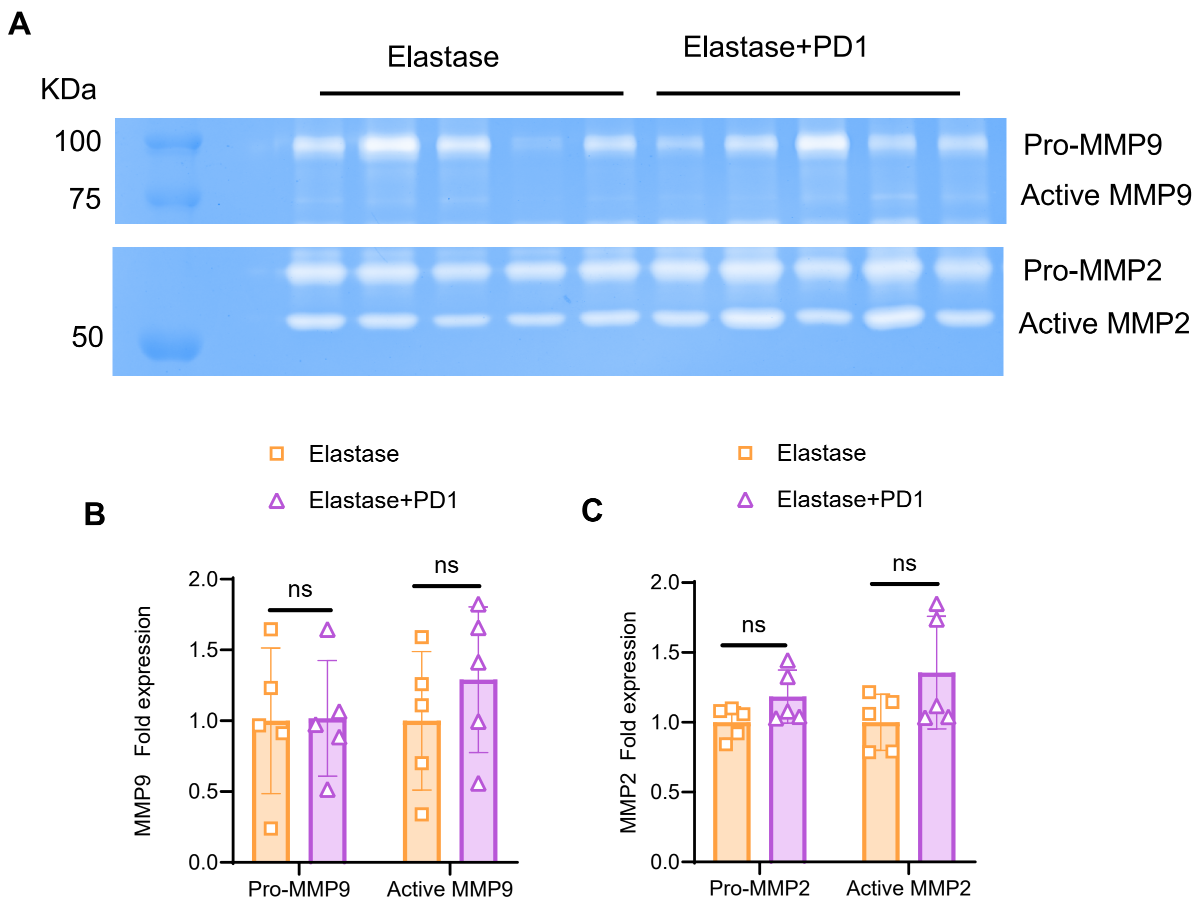


**Supplementary Figure S4. A.** Gelatin zymography of murine aortic tissue in GPR37$^{-/-}$ mice and **(B-C)** quantification of optical density (O.D.) shows no significant change in levels of MMP9 and MMP2 expressions, after PD1 treatment compared to respective elastase-treated groups. ns, not significant; n=5/group; ns, not significant.

**Supplementary Figure S5**

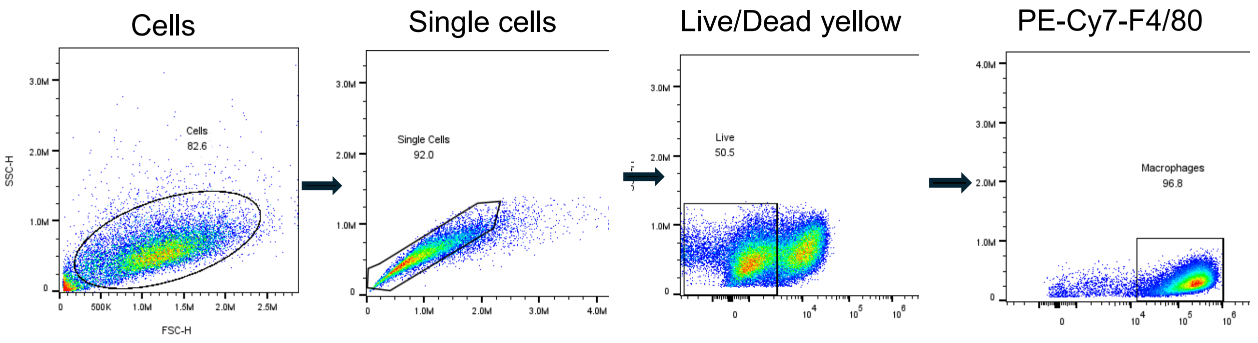


**Supplementary Fig. S5**. Gating strategy for flow cytometry analysis of efferocytosis for *in vitro* cultures of macrophages and SMCs.

**Supplementary Figure S6**

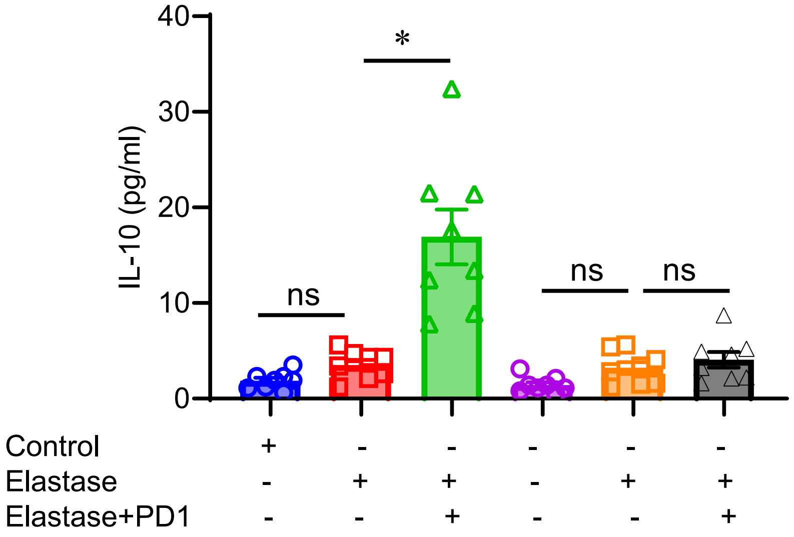


**Supplementary Fig. S6.** Purified macrophages from WT and GPR37$^{-/-}$ mice were exposed to transient elastase-treatment and treated with/without PD1 treatment. IL-10 expression was significantly increased in PD1-treated culture supernatants in WT but not GPR37$^{-/-}$ macrophages compared to respective untreated controls. *p<0.001; ns, not significant; n=8/group.
